# The West African lungfish secretes a living cocoon during aestivation with uncertain antimicrobial function

**DOI:** 10.1101/2024.07.05.602297

**Authors:** M. Fernanda Palominos, Rangarajan Bharadwaj, Charles Tralka, Kenneth Trang, David Aka, Mariam Alami, Dominique Andrews, Ben I. Bartlett, Chloe Golde, Joseph Liu, Maya Le-Pedroza, Robert Perrot, Blanca Seiter, Claudia Sparrow, Michael Shapira, Christopher H. Martin

## Abstract

One of the most exceptional adaptations to extreme drought is found in the sister group to tetrapods, the lungfishes (Dipnoi), which can aestivate inside a mucus cocoon for multiple years at reduced metabolic rates with complete cessation of ingestion and excretion. However, the function of the cocoon tissue is not fully understood. Here we developed a new more natural laboratory protocol for inducing aestivation in the West African lungfish, *Protopterus annectens,* and investigated the structure and function of the cocoon. We used electron microscopy and imaging of live tissue-stains to confirm that the inner and outer layers of the paper-thin cocoon are composed primarily of living cells. However, we also repeatedly observed extensive bacterial and fungal growth covering the cocoon and found no evidence of anti-microbial activity in vitro against *E. coli* for the cocoon tissue in this species. This classroom discovery-based research, performed during a course-based undergraduate research experience course (CURE), provides a robust laboratory protocol for investigating aestivation and calls into the question the function of this bizarre vertebrate adaptation.

## Introduction

Evolutionary novelties provide fascinating subjects for engagement with science, tests of evolutionary theory, and case studies for mapping the genetic basis of human diseases (Streelman et al. 2007; Moczek 2008; Shubin et al. 2009; Powder and Albertson 2016; Davis et al. 2019). One set of examples are provided by ephemeral and intromittent aquatic habitats, which have selected for the convergent evolution of aestivation in a diverse group of aquatic and semi-aquatic vertebrates (reviewed in (Glass et al. 2009; Secor and Lignot 2010; Lajus and Alekseev 2019). This includes spadefoot toads (Zamora-Camacho et al. 2019; Calabrese and Pfennig 2023; Chen et al. 2023), African clawed frogs (Childers 2014), turtles (Ligon and Stone 2003), amphiumas (Smith and Secor 2017), and the Australian salamanderfish (Ogston et al. 2016) among many other species known to aestivate. One of the most exceptional examples of aestivation are found in the South American (*Lepidosiren paradoxa*) and African lungfishes (*Protopterus* spp.), known to withstand multi-year droughts as adults curled inside a mucus cocoon (Smith 1931; Janssens 1964; Reno et al. 1972). Dipnoi are the sister group to all tetrapods, representing over 400 million years of independent evolutionary history and potentially novel strategies for surviving droughts over this immense timespan (Criswell 2015). As water levels fall, lungfishes remain in their muddy burrows and shed additional mucus to form a thin papery cocoon which was originally thought to be reminiscent of dried leaves (Smith 1931). The only opening in the cocoon is to their mouth for respiration which is maintained through a narrow passage to the surface from their muddy burrow. During this time they cease all feeding and excretion, shift from the production of ammonia to urea, and substantially drop their metabolic and respiratory rates in a state of torpor (Chew et al. 2004; Loong et al. 2008; Hiong et al. 2013; Chew et al. 2015). They can remain in this state for at least several years, losing over 10% of their body mass until the rains return, when they begin normal body movement and foraging activities within a day (Janssens 1964; Fishman et al. 1986; Greenwood 1986; Fishman et al. 1992; Glass et al. 2009; Chew et al. 2015).

It was recently reported that the slender African lungfish (*Protopterus dolloi*) secretes a cocoon composed of living tissue with antimicrobial properties drawing from large reservoirs of granulocytes in its organs (Heimroth et al. 2021; Salinas et al. 2023). Cocoons were examined after ten days following food restriction and antimicrobial function was inferred from the presence of extracellular protein traps, high levels of beta defensin expression, and potentially new skin toxins (Tacchi et al. 2015; Heimroth et al. 2018, 2021; DeMmon et al. 2022; Casadei and Salinas 2023; Salinas et al. 2023). However, many aspects of cocoon function are still unknown, particularly after longer time periods in aestivation, between inner and outer layers of the cocoon, and in additional lungfish species besides *P. dolloi*.

Here we first developed a new laboratory protocol that does not involve food restriction for inducing aestivation and cocoon formation in the West African lungfish (*P. annectens*) and includes the addition of loam-rich wet soil to better recreate the natural conditions surrounding aestivation in this species and avoid unnecessarily restricting food. We confirmed that the paper-thin cocoon tissue of the West African lungfish is composed predominantly of living cells on both the inner and outer surfaces using fluorescent nuclear and cell integrity staining, consistent with its potential role in immune function. However, we found no evidence of antimicrobial activity of cocoon tissue using standard *E. coli* growth and inhibition assays. Overall, we find the West African lungfish to be a fascinating laboratory model for course-based undergraduate research experience (CURE) courses, during which the aestivation protocol and results reported here were pioneered by undergraduate student coauthors over the past three years.

## Methods

Wild-caught West African lungfish (*P. annectens*) imported from Nigeria (*n* = 3) were acquired from U.S. commercial retailers in 2021 and 2022. Adult fish (30 cm in length) were housed in flow-through 400-liter partitioned acrylic tanks at 23-27° C, pH 8, with a 12:12 photoperiod under artificial light following standard husbandry conditions for other freshwater fishes in the lab (Martin 2012; Martin et al. 2019; Palominos et al. 2023). Fish were fed every other day with primarily commercial pellets (New Life Spectrum and Hikari) supplemented with occasional frozen bloodworms (Hikari) or live feeder fish. Fish were housed in the laboratory for at least two months before use in any experiments.

### Aestivation protocol

In order to monitor aestivation visually without disturbing the fish, we developed a new experimental procedure to induce aestivation in the lab (see also (Delaney et al. 1974; Ip et al. 2005). Terrestrial loam-rich soil with minimal organic matter was collected from the UC Berkeley campus. Because the lungfish were not sterile, we did not sterilize the soil but took care to avoid collecting soil from aquatic habitats to avoid contamination or potential pathogen exposure. An approximately 2 cm layer of soil covered with tank water to 2 cm depth was placed in a clean 40-liter aquarium under a photoperiod of 12:12. Each lungfish was placed directly into its own tank without any preceding period of starvation at 23-25° C ambient air temperatures, taking care to maintain high levels of humidity with a tight-fitting lid (but not completely airtight) and mud along the bottom of the tank. Over the course of a few days, lungfish progressively suspended movement and secreted an increased amount of mucus from their opercula as water evaporated naturally from each tank, leaving only a layer of mud (Fig. 1). Within approximately 1-2 weeks, depending on the amount of residual water in each tank, the lungfish cocoon dried around each animal alongside the hardened mud after most residual water evaporated. Animals were monitored daily for any signs of desiccation, but no additional food or water was provided during the aestivation period, except a few milliliters of water added to the surrounding hardened mud as needed (approximately weekly) to maintain humidity at 80-100%. One animal responded with an audible ‘barking’ vocalization in response to light touch but did not respond in later aestivation trials (initially reported by (Smith 1931)). This was a vocalization in response to touch and distinct from respiration which could be observed at the opercula for approximately the first week before the cocoon hardened around the lungfish. Initial pilot experimental trials were unsuccessful in inducing aestivation if a deep layer of mud (0.3 m) was provided, which may substantially prolong the time needed for cocoon formation.

**Fig. 1.**
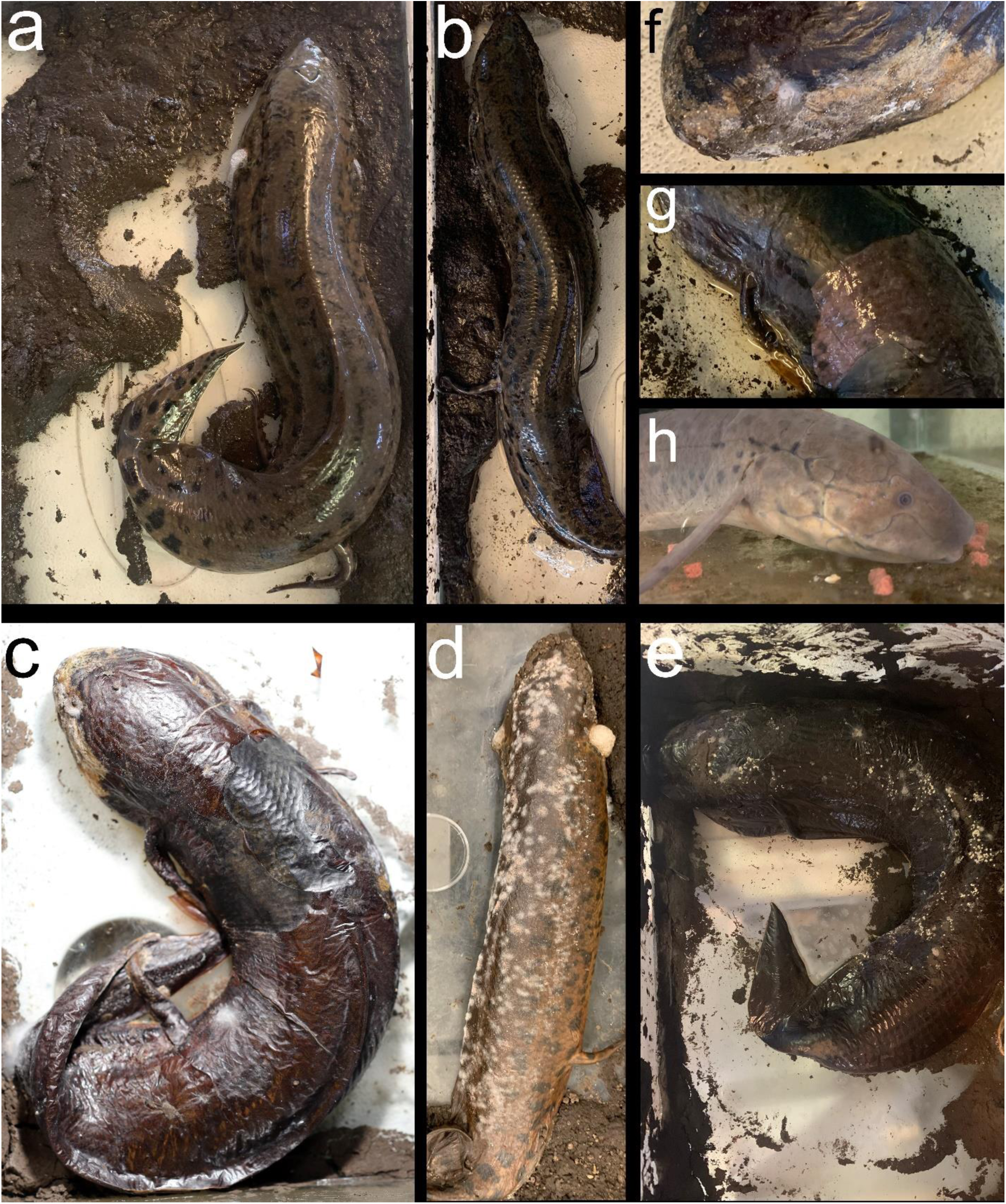
West African lungfish in various states of aestivation and recovery. a-b) Newly added to 40-liter terrarium with mud layer. c-f) After multiple weeks and months in aestivation. Note the papery thin cocoon covered in mold and fungal colonies in some areas. g) Removal of cocoon following addition of tank water after approximately one hour. h) One week post recovery after aestivation period of three months, readily feeding on commercial pellet food.

Animals were maintained in a state of aestivation for up to three months with no ill effects as long as humidity levels remained high within each covered tank (Fig. 1). Lungfish could be recovered from their state of aestivation by simply adding dechlorinated water to the aestivation tank to a depth of approximately 2-3 cm. After approximately one hour, the cocoon tissue softened, sloughed off, and the animals began to respond slowly to gentle touch. Cocoon tissue could either be peeled away or left attached. Lungfish were returned to their home tanks with a reduced water level to allow easy access to the water surface for respiration and were provided with a small amount of food. Within 24 hours, animals were eating normally and within a few days had completely recovered typical levels of movement and response to stimuli. Repeated aestivation trials on the same animal were possible with a recovery period of only one week. However, all samples were collected from animals that had recovered from aestivation in laboratory aquaria for at least three months before repeating an aestivation trial.

### Sampling cocoon tissue

After at least two weeks in aestivation, the mucus layer hardened into a thin papery shell around each fish (Fig. 1). Samples of this tissue after approximately 1 month in aestivation were removed by carefully puncturing the cocoon with forceps at an angle parallel to the surface of the fish to avoid damaging the inner tissue. The cocoon tissue layer could then be peeled off for imaging, staining, and antimicrobial assays. We noted that cocoon tissue reformed around these sampling areas after approximately one week (darkened central dorsal patches visible in Fig. 1c). Cocoon tissue was always sampled fresh from the aestivating animal before procedures and never preserved before use.

### Scanning electron microscopy

Fresh cocoon tissue samples were prepared for electron microscopy imaging using standard protocols by the Electron Microscope Lab at UC Berkeley. Samples were fixed in 2% glutaraldehyde in 0.1M sodium cacodylate buffer for 1-2 hours, rinsed in 0.1M sodium cacodylate buffer, and then post-fixed in 1% osmium tetroxide in 0.1M sodium cacodylate buffer. After three rinses with 0.1M sodium cacodylate buffer, samples were dehydrated in a stepped series of ethanol washes. Following critical point drying, each sample was cut into at least 2 parts and placed upwards and downwards onto the SEM stub, resulting in images taken from both the outer and inner surface of the cocoon. Samples were mounted onto stubs by using conductive carbon tape or silver paint and imaged on a Zeiss Crossbeam 550 FIB-SEM.

### Cocoon tissue staining

Fresh cocoon tissue sampled from two aestivating lungfish was double-stained with propidium iodide (Thermo Fisher Scientific) to detect newly dead cells from the damage to the integrity of the cell membrane and DAPI (Sigma-Aldrich) to label cell nuclei. Stains were conducted at two different timepoints after initiating aestivation trials: one month post and three months post aestivation. Briefly, the dissected pieces of the cocoon were mounted on positively charged (X) Poly-L lysine (Sigma-Aldrich) coated slides and stained for 15 min covered from light with a staining solution containing propidium iodide and DAPI, respectively. We stained and imaged both the inside and outside of the cocoon, sampled from the dorsal medial surface of the fish.

### Antimicrobial assays

Fresh cocoon tissue was sampled from two aestivating lungfish at two different timepoints after initiating aestivation trials: two weeks post and six weeks post. *E. coli* (OP50 strain originally obtained from the Caenorhabditis Genome Center (Brenner 1974)) was prepared by inoculating a 1L flask of liquid Luria Broth (LB; 10g Tryptone, 5g Yeast Extract, 5g NaCl, to 1L H2O, autoclaved) from a single streaked colony, and then incubated at 25°C overnight to saturation, then concentrated tenfold via centrifugation, and lastly stored at 4°C for up to a week prior to use. Either 24 hours before exposure to tissue to test for bactericidal compounds or a few hours before tissue exposure to test for bacterial growth inhibitors, 500 µL of 10X concentrated OP50 was inoculated and sterilely spread on LB agar plates (LB + 15g agar). Dorsal sections of cocoon tissue were placed on each plate (Fig. 4) and incubated at room temperature to carefully monitor bacterial growth over the following seven days. We incubated at lower than optimal temperatures to carefully track bacterial growth progress over multiple days. We tested a total of 16 plates at both inoculation conditions. We also exposed a set of control plates that were not inoculated with *E. coli* (*n* = 4). In all cases, we examined plates for evidence of bactericidal or growth inhibition around the cocoon tissues after 1, 3, and 7 days post-exposure.

## Results

We successfully pioneered a new and efficient laboratory protocol for inducing aestivation in the West African lungfish. We confirmed that repeated aestivation trails and repeated sampling of tissue from the same region were possible. In all aestivating animals (*n* = 3) across three sets of trials over three years (2021, 2022, and 2023), we observed extensive growth of mold on the surface of cocoons and in the surrounding mud, usually within two to three weeks after initiating aestivation. This growth did not appear to impact the subsequent health of the animal but does call into question the anti-microbial properties of the cocoon tissue during early stages of aestivation. However, no mold growth was observed on the inner surface of the fish after removing cocoon tissue, so the cocoon may still be providing a barrier to microbial entry. Scanning electron micrographs confirmed the uneven, semi-porous structure of the outer surface of the cocoon (Fig. 2), which was also covered with bacteria, fungal spores, and fungal hyphae (Fig. 3).

**Fig. 2.**
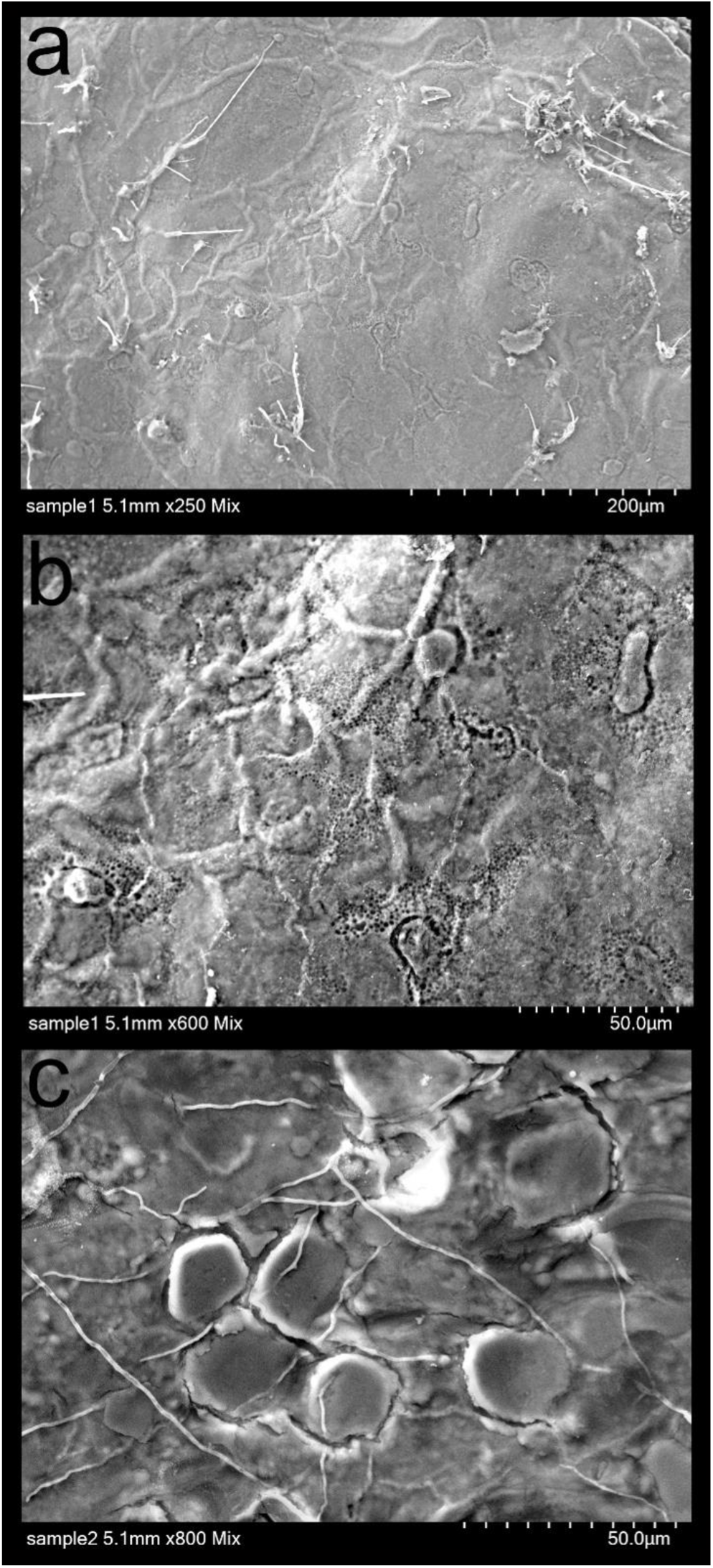
SEM images of the outer surface of lungfish cocoons from two different aestivating animals. a-c) Note the cocoon filaments and in some cases porous texture of the surface of the cocoon.

**Fig. 3.**
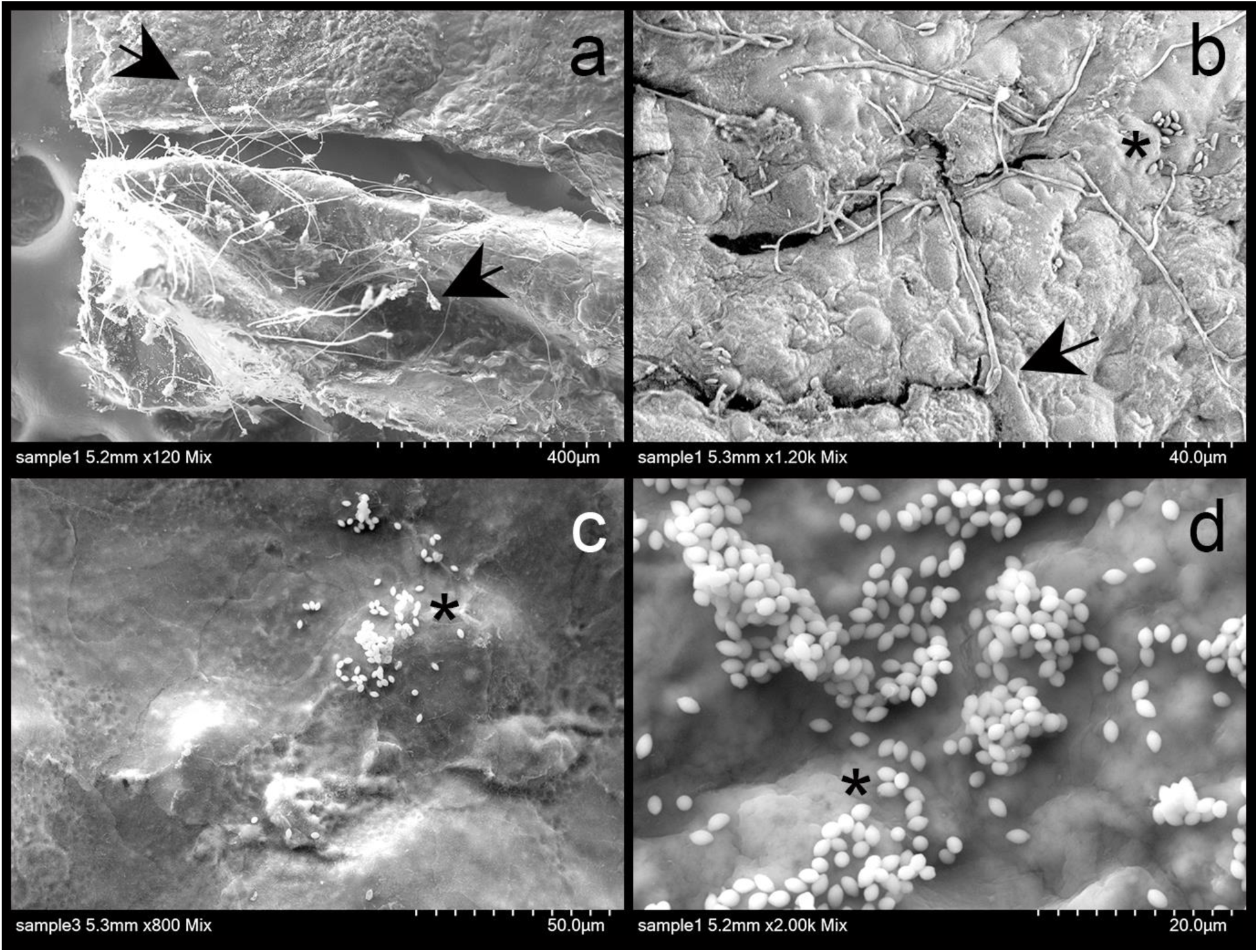
SEM images of the outer surface of lungfish cocoon from two aestivating individuals. a-b) Fungal hyphae. c-d) Aggregation of possible yeast. Arrows pointing to hyphae and asterisks to spores.

Antimicrobial assays using *E. coli* inoculated LB plates provided no evidence of bactericidal or bacterial growth inhibition of cocoon tissue samples from multiple animals at two different timepoints during aestivation trials in 2023. In all cases, a ‘halo’ of reduced growth was not observed surrounding tissue samples placed in the center of each plate, whether exposed to tissue 24-hours post-inoculation or a few hours after inoculation (Fig. 4). Instead, increased bacterial growth was observed around each sample over several days post exposure. To test if this was *E. coli* growth or another microbe, we also examined sterile LB plates exposed to cocoon tissue and observed the same increased level of microbial growth around each sample, potentially due to the bacterial communities already present on the lungfish cocoon.

**Fig. 4.**
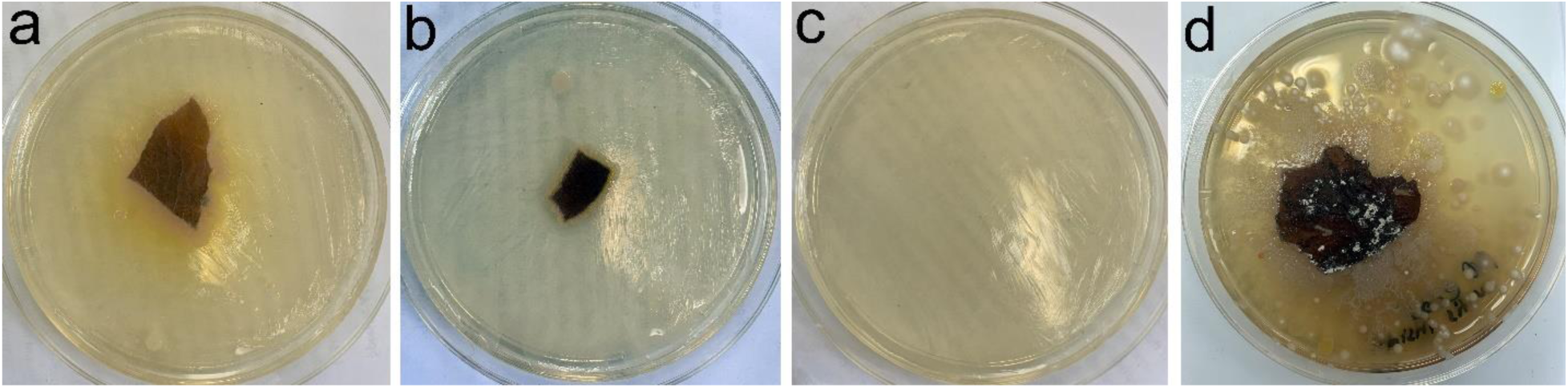
Antimicrobial assays using LB plates inoculated with E. coli. a-b) Lungfish tissue removed after two months of aestivation shows no evidence of antimicrobial activity against E. coli growth rings surrounding tissue relative to normal E. coli growth in c) control plate. d) LB plate without *E. coli* inoculation shows extensive microbial growth surrounding cocoon tissue.

Nonetheless, double-staining of both the wet inside layer and dry outside layer of the cocoon from multiple animals at two timepoints during aestivation indicated that most of the cells on the inner and outer layer of the lungfish cocoon are alive, consistent with previous results in a different species of lungfish, *P. dolloi* (Heimroth et al. 2021). Propidium iodide staining indicated that dead cells within the cocoon tissue occurred relatively infrequently on both inner and outer layers (Fig. 5, arrows, Fig. 5c, inset). In contrast, DAPI cell nuclei staining indicated dense and regular spacing of cells across both the inner and outer cocoon sections examined (Fig. 5); however, besides the regularly-spaced nuclei, we also observed several neutrophil-like shaped nuclei (Fig. 5d-f, arrowheads, Fig. 5f, inset) in the outer region of the cocoon of one of the aestivating *P. annectens*. This is also in accordance with the reported infiltration of granulocytic immune cells from the aestivating lungfish to the cocoon (Heimroth et al. 2021), alongside what looks like neutrophilic extracellular traps (Fig. 5d-f, asterisks)

**Fig. 5.**
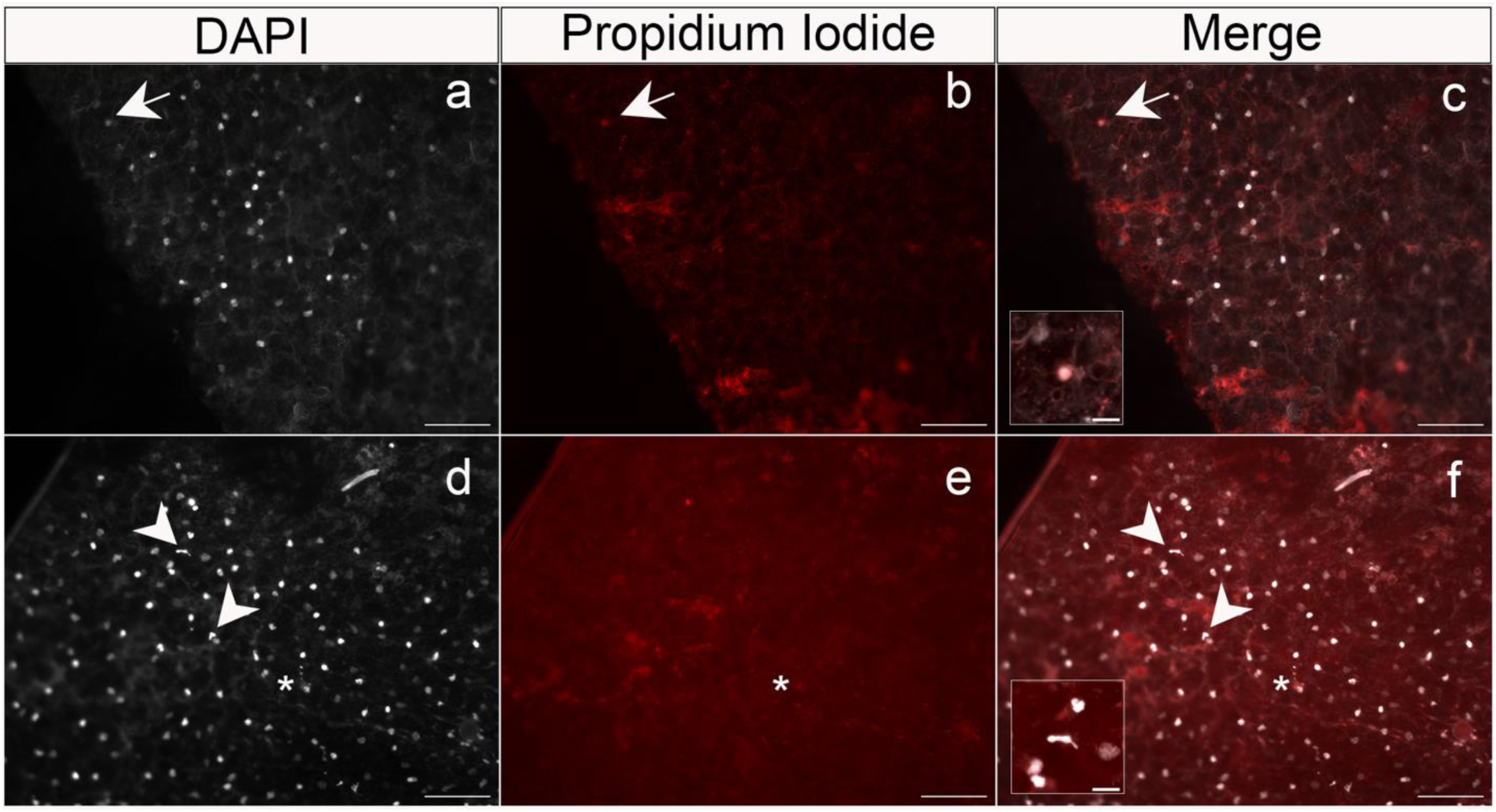
Double-staining of the inner region from two lungfish cocoon tissue samples with DAPI (left column) and propidium iodine (PI, middle column). Images were taken on a Axio Imager M2 fluorescent microscope. Merged images are shown in c) and f). Tissues were sampled fresh from two different *P. annectens* aestivating individuals for two weeks. a) Note the greatly increased number of DAPI-stained nuclei present in the left column relative to only a b) single PI positive nuclei (arrows), pointing out a unique cell in the later stages of cell death among all DAPI-positive PI-negative nuclei (c, inset). d-f) Arrowheads indicate neutrophil-like shaped nuclei, one of them magnified in f). Scale: 100 µm for all pictures, and 20µm for the insets. Asterisks indicate possible neutrophil extracellular traps.

## Discussion

We developed a new, efficient and more natural laboratory protocol for inducing aestivation in the West African lungfish *P. annectens* through an undergraduate course-based undergraduate research experience (CURE) class *Ichthyology: an introduction to the scientific process through the study of fishes.* This format has resulted in many successful research projects in which students form their own hypotheses and test their ideas during the lab period over a semester, resulting in both student-led publications (Zeng and Martin 2017; Davis et al. 2019; St John et al. 2020; Tan et al. 2023) and contributions to larger research projects in the lab (St. John and Martin 2019; St John et al. 2019; St. John et al. 2020, 2021; Richards et al. 2021; Galvez et al. 2022). We discovered that both the inner wet and outer dry layer of the mucus cocoon is composed predominantly of living cells (Fig. 4), consistent with earlier work showing that the cocoon tissue is alive in a different species of African lungfish (Heimroth et al. 2021). Moreover, we also pioneered a working protocol to stain and quantify live and dead cells on aestivating lungfish cocoon tissue samples.

We observed an alarming amount of mold and microbial growth on this living cocoon over repetitive aestivation trials (Figs. 1-2) in a non-sterile environment, similar to conditions in the wild during aestivation. Nonetheless, animal health does not appear to be affected by microbial growth on the outer layer of the cocoon which may prevent microbes from reaching the inner cocoon layer next to the aestivating lungfish’s skin (Fig. 1d). Recovery time may be needed between induced aestivation trials for the lungfish to be able to produce a functional healthy cocoon that will protect them from the outer desiccating and unsterile environment.

We found no evidence of antimicrobial activity against *E. coli* growth or bactericidal activity by the cocoon tissue itself. However, additional assays against a wider range of microbial taxa in both liquid media and plates are needed to determine whether this tissue plays a role in immune function. It is also unclear whether recruitment from granulocyte reservoirs in forming the cocoon tissue necessarily leads to immune function by this tissue over the full span of aestivation in natural conditions (Ip et al. 2005; Heimroth et al. 2021).

In favor of the immune function hypothesis for the lungfish cocoon, we found neutrophil-like shaped nuclei and what looks like neutrophil extracellular traps in the inner wet region of *P. annectens* cocoons (Fig. 5); however, we did not find that the cocoon itself was mainly composed of granulocytes or other innate immune cells. Instead, the cocoon shows a high degree of cell nuclei organization suggesting that it is a complex living organized tissue which harbors a plethora of cell types that have different organizational requirements. Overall, our results highlight the still unknown function of an organized external living tissue in Dipnoi.

Alternatively, the cocoon may function to regulate metabolism and air exchange with the aestivating lungfish, which largely ceases to breathe and stops all ingestion and excretion of waste during this period (Chew et al. 2004, 2015; Ip et al. 2005). Furthermore, it remains unknown how the living cells within the cocoon are nourished or supplied with oxygen through passive diffusion or some sort of additional circulatory network.

Ultimately, no other aestivating vertebrate animals are known to secrete a living cellular matrix during aestivation to our knowledge (Glass et al. 2009; Secor and Lignot 2010; Lajus and Alekseev 2019). However, given that this was reported only recently in lungfish, likely few if any other studies have investigated the mucus layers surrounding other aestivating vertebrate groups which were previously assumed to be dried, desiccated mucus or interlocked layers of proteins (Glass et al. 2009; Secor and Lignot 2010; Storey and Storey 2012). The investigation of extraordinary evolutionary novelties can often lead in unexpected directions and is increasingly urgent during this period of rapid biodiversity loss during the Anthropocene (Moyle and Leidy 1992; Darwall and Freyhof 2016; Johnson et al. 2017).

## Acknowledgements

We thank the University of California, Berkeley, NSF CAREER 1749764, NIH NIDCR 5R01DE027052-02, the Berkeley Collegium Fund, and a Berkeley Discovery-based learning grant for funding to CHM. We also thank Danielle Jorgens at the Electron Microscope Lab at UC Berkeley for quickly processing our samples in collaboration with undergraduate researchers. All animal care protocols (AUP-2021-02-14062-1 and AUP-2021-07-14515) were approved by the University of California, Berkeley Animal Care and Use committees.

## Data Accessibility

Data will be deposited to Dryad digital repository.

